# Africa’s Genetic Heritage in the Digital Era:DSI Governance Gaps, Multi-Forum Benefit-Sharing Architecture, and Strategic Priorities for COP17

**DOI:** 10.64898/2026.05.18.725948

**Authors:** Yves Shema, Suwilanji Sinyangwe, Fredrick Abu Ayodele

## Abstract

**Background:** Africa is among the most biologically rich continents on Earth for non-human life, yet structurally excluded from the benefit-sharing framework that the Convention on Biological Diversity (CBD) was designed to create. Digital sequence information (DSI), nucleotide and protein sequence data derived from non-human organisms including plants, animals, fungi, microbiota, pathogens, and human microbiome-associated micro-organisms, underpins industries generating over USD 1.56 trillion in annual revenue. DSI governance operates across multiple international forums simultaneously: the CBD’s Cali Fund (operationalised in 2024) is the primary mechanism, but the International Treaty on Plant Genetic Resources for Food and Agriculture (ITPGRFA), the World Health Organization (WHO) Pandemic Instrument framework, the Biodiversity Beyond National Jurisdiction (BBNJ) Agreement, and the new World Intellectual Property Organization (WIPO) Treaty on Intellectual Property, Genetic Resources and Associated Traditional Knowledge each govern distinct categories of DSI. Africa’s claims to equitable benefit-sharing rest primarily on its extraordinary non-human biodiversity, but the governance architecture is fragmented, voluntary, and structurally ill-equipped to deliver returns to the communities and nations that are the custodians of these resources.

**Objectives:** This study (i) characterises Africa’s non-human biodiversity endowment and its underrepresentation in global DSI databases; (ii) maps access and benefit-sharing (ABS) legislative readiness across 54 African Union (AU) member states, accounting for pre-DSI legal frameworks and national heterogeneity; (iii) models commercial trajectories and Cali Fund benefit-sharing scenarios; (iv) synthesises the multi-forum DSI governance architecture; and (v) elicits, through a structured key-informant exercise, the governance preferences, disbursement priorities, and benefit-sharing modalities of 25 CBD delegates and experts consulted at SBSTTA-28/SBI-7 (Nairobi, July-August 2026).

**Methods:** A structured evidence synthesis was conducted following PRISMA 2020 reporting elements. Literature was searched across PubMed, Scopus, Web of Science, and Google Scholar, supplemented by official CBD, FAO, IUCN, UNEP, WHO, WIPO, and BBNJ repositories. The search used expanded terminology beyond “DSI” to include “genetic sequence information,” “biological sequence data,” and country-specific ABS law terms to avoid false negatives from narrower DSI-restricted searches. Quantitative Cali Fund scenario modelling used the UNEP-WCMC/KPMG (2024) non-human DSI sector revenue baseline. To complement the synthesis, a structured expert elicitation was undertaken with 25 purposively selected key informants (CBD Party delegates, IPLC representatives, scientific experts, and ABS focal points) at SBSTTA-28/SBI-7 (Nairobi, July-August 2026). Consistent with the logic of expert elicitation and key-informant sampling, these data are treated as qualitative, position-mapping evidence indicative of negotiator and expert perspectives, not as a statistically representative sample.

**Results:** Africa’s non-human biodiversity endowment is exceptional: approximately 25% of global terrestrial species, 9 of 36 biodiversity hotspots, and the world’s second-largest tropical forest. Yet African non-human taxa are severely undersequenced in International Nucleotide Sequence Database Collaboration (INSDC) databases, reflecting both a collection gap and a metadata attribution problem. Zero AU member states have enacted ABS legislation explicitly covering non-human DSI; however, several states have pre-DSI genetic resources laws (e.g., Namibia’s 2017 legislation) that may provide partial coverage under expanded interpretation. Among the 25 informants consulted (representing 14 countries), a clear majority (20 of 25) favoured mandatory contribution models; informants ranked centre of origin and documented material use above raw biodiversity share as disbursement factors, and identified technology transfer and capacity building as the most valuable benefit forms alongside monetary payments. Twenty-two of 25 informants indicated FAO ITPGRFA should govern or coordinate on plant DSI; 20 of 25 flagged the WHO framework for pathogen sequences. These convergent expert views corroborate, rather than establish on their own, the conclusions drawn from the literature synthesis.

**Conclusions:** Africa’s modelled allocation share under the Cali Fund ranges from USD 312 million to USD 5.83 billion annually depending on levy rate and biodiversity weight. Realising this requires urgent legislative reform, a shift from voluntary to mandatory contribution mechanisms, multi-forum coordination, and direct IPLC engagement. The domesticated species and legacy DSI complications require a multi-criteria disbursement formula that goes beyond simple biodiversity share. Because the elicitation is a purposive, hypothesis-generating exercise rather than a representative survey, a research agenda for representative, field-based primary data collection is identified to convert these indicative findings into a quantified evidence base.

## 1. Introduction

The governance of genetic resources and the digital sequences derived from them sits at the intersection of international law, biotechnology, conservation science, and global equity. Africa is among the world’s most biologically rich continents for non-human life, yet structurally excluded from the benefit-sharing framework that the CBD was designed to create. This paradox has deepened with the digitisation of biology: once a genetic sequence from an African plant, pathogen, or soil microbe is uploaded to an open-access database, the source country currently has no mechanism to track its commercial use or assert a benefit-sharing claim.

Digital sequence information (DSI), nucleotide and protein sequence data derived from non-human organisms and deposited in public databases, is now the primary substrate for innovation in pharmaceuticals, agricultural biotechnology, cosmetics, and diagnostics. The non-human DSI-dependent industries generated over USD 1.56 trillion in annual revenue in 2024.[1] A critical legal distinction must be stated at the outset: the human exception to CBD sovereign rights does not extend to pathogens, micro-organisms associated with the human microbiome, or any other non-human organism. States retain full sovereign rights over access to these resources and to DSI derived from them, regardless of their association with human health. Human genetic resources (i.e., the human genome itself) are governed separately under the WHO framework and bioethics instruments; this study addresses them only as a secondary infrastructure indicator, as detailed in Section 4.2.2.

DSI governance is not a single-forum matter. While the CBD’s Cali Fund is the primary mechanism for non-human biodiversity-based DSI, parallel international processes govern specific categories: the FAO International Treaty on Plant Genetic Resources for Food and Agriculture (ITPGRFA) for plant DSI in food and agriculture; the WHO Pandemic Instrument for pathogen sequences; the BBNJ Agreement for marine DSI beyond national jurisdiction; and the new WIPO Treaty on Intellectual Property, Genetic Resources and Associated Traditional Knowledge for patent-linked DSI disclosure. A comprehensive analysis of Africa’s strategic position must account for this multi-forum architecture.

This study makes five contributions: (i) it characterises Africa’s non-human biodiversity endowment as the primary basis for Cali Fund allocation claims; (ii) it maps Africa’s ABS legislative readiness across 54 AU states, explicitly accounting for pre-DSI legal frameworks and national heterogeneity; (iii) it models Cali Fund economic trajectories under alternative parametric assumptions; (iv) it synthesises the multi-forum DSI governance architecture relevant to Africa; and (v) it incorporates a structured expert elicitation of 25 CBD delegates and experts at the SBSTTA-28/SBI-7 meetings in Nairobi (July-August 2026), constituting the first documented account of negotiator and expert preferences on DSI governance and benefit-sharing priorities. Consistent with established elicitation practice, this component is used to map and triangulate expert positions rather than to estimate population parameters; a research-gap and future-directions section identifies the representative primary-data agenda needed to make the analysis conclusive.

## 2. Conceptual and Regulatory Background

### 2.1 Defining Non-Human DSI and the Legal Scope of CBD Coverage

The term “digital sequence information” was introduced into CBD negotiations at COP13 (Cancún, 2016) through Decision XIII/16. In the context of the CBD and the Cali Fund, DSI encompasses nucleotide and protein sequence data from non-human organisms, plants, animals, fungi, microbiota, and pathogens. As of 2024, GenBank, maintained by the US National Center for Biotechnology Information (NCBI) and the flagship of the International Nucleotide Sequence Database Collaboration (INSDC), contains 25 trillion base pairs from over 3.7 billion sequences for 557,000 formally described species.

The INSDC’s 2023 update required collection-location and collection-date metadata for all new BioSample registrations, the most significant step toward non-human DSI traceability; however, billions of legacy records carry no country-of-origin metadata.[2] This legacy data problem is a central governance challenge: sequences collected before current frameworks carry no retroactive Nagoya obligations, yet they are commercially exploited in perpetuity. The Cali Fund’s multilateral model was specifically designed to address this legacy data problem by creating a collective contribution mechanism that does not require transaction-by-transaction traceability.

An important distinction raised at the DSI side event (28 July 2026) is between ‘old DSI’ (sequences derived from biological material collected before the Nagoya Protocol entered into force in 2014) and ‘new DSI’ (sequences from post-2014 collections). Legacy sequences carry no retroactive Nagoya obligation; post-2014 sequences should in principle be traceable to their access agreements. This temporal divide affects how Africa frames its allocation claims: for legacy sequences, the multilateral fund model is the only viable mechanism; for new sequences, bilateral and multilateral mechanisms may both apply.

### 2.2 The Multi-Forum DSI Governance Architecture

**Table 1.** Governance of Digital Sequence Information (DSI) by Category and Relevance for Africa.

| DSI Category | Governing Instrument | Mechanism | Relevance for Africa |
| --- | --- | --- | --- |
| Wild flora, fauna, fungi, microbiota | CBD / Nagoya Protocol / Cali Fund | Multilateral fund; voluntary contributions | Primary basis for Africa's allocation claim. All wild African taxa. |
| Plant genetic resources for food & agriculture (domesticated crops) | FAO ITPGRFA (Annex 1 crops) / CBD for non-Annex crops | Multilateral System; Standard Material Transfer Agreement | Complex: many staple crops (wheat, rice, maize) originate outside Africa. Africa must engage in ITPGRFA reform. |
| Pathogen sequences (viruses, bacteria causing human disease) | WHO Pandemic Instrument / CBD (non-human status confirmed) | Pandemic Influenza Preparedness (PIP) Framework model; in negotiation | Africa holds significant pathogen diversity. WHO instrument must benefit African public health institutions. |
| Marine DSI beyond national jurisdiction | BBNJ Agreement (2023) | Multilateral benefit-sharing; identifier model at collection | Africa has extensive EEZ. BBNJ's sample identifier model is a provenance-tracking template for terrestrial DSI. |
| Patent-linked DSI (innovations based on genetic resources) | WIPO Treaty on IP, Genetic Resources & Associated TK (2024) | Patent applicants must disclose country of origin of genetic resources | New enforcement angle: creates paper trail linking commercial IP to African source material. |

*Note.* DSI = digital sequence information; CBD = Convention on Biological Diversity; FAO ITPGRFA = International Treaty on Plant Genetic Resources for Food and Agriculture; PIP = Pandemic Influenza Preparedness; BBNJ = Biodiversity Beyond National Jurisdiction; WIPO = World Intellectual Property Organization; TK = traditional knowledge; EEZ = exclusive economic zone.
*Sources:* CBD Decision 16/2 [7]; FAO ITPGRFA [8]; BBNJ Agreement [9]; WIPO Treaty on IP/GR/TK [10]; WHO PIP Framework [11].

The BBNJ Agreement’s model of attaching unique identifiers to biological samples at the point of collection, so that the identifier follows the sequence into any database, is particularly relevant as a provenance-tracking template for terrestrial DSI. This model is technically feasible, as demonstrated by the BBNJ negotiations, and the side event (28 July 2026) recommended evaluating whether it can be adapted for African terrestrial biodiversity programmes.[9]

The WIPO Treaty on Intellectual Property, Genetic Resources and Associated Traditional Knowledge (adopted May 2024) requires patent applicants to disclose the country of origin of genetic resources used in inventions. This creates a new traceability mechanism that directly links commercial patent applications to source countries, generating a legal paper trail that could significantly strengthen Africa’s DSI traceability claims.[10]

### 2.3 CBD Regulatory Architecture and the Cali Fund

The CBD (1992, 196 Parties) establishes three co-equal objectives: conservation of biological diversity; sustainable use; and fair and equitable benefit-sharing from genetic resources. The Nagoya Protocol (adopted 2010, entered into force 2014, 142 countries) operationalises the third objective through a bilateral ABS framework governing non-human genetic resources.[4,5]

The Kunming-Montreal Global Biodiversity Framework (KMGBF), adopted at COP15 in December 2022, sets 23 global targets for 2030. Target 13 addresses non-human DSI directly, and Goal C stipulates fair and equitable sharing of benefits from the utilisation of non-human genetic resources and DSI. CBD Decision 16/2 (November 2024; authoritative text: CBD/COP/16/L.19) operationalised the Cali Fund with indicative, voluntary contribution rates of 1% of profits or 0.1% of revenue. Final contribution rates are to be determined at COP17 (Yerevan, Armenia, October 2026).[7,12,13]

### 2.4 The Domesticated Species Complication and Africa’s Allocation Claim

Africa’s 9/36 global biodiversity hotspot share (25%) provides a defensible upper bound for allocation from wild biodiversity. However, a significant proportion of the commercially valuable DSI concerns domesticated crops (wheat, rice, maize, soybean) and livestock whose primary centres of genetic origin are in the Fertile Crescent, East Asia, or Mesoamerica, not Africa. The FAO ITPGRFA governs these resources through a separate multilateral system.

For Africa’s wild biodiversity, endemic plant species, wildlife, marine fauna, soil microbiomes, and the Congo Basin ecosystem, the Cali Fund biodiversity-share allocation model is most appropriate and most defensible. Africa’s allocation claim is strongest for wild, non-domesticated organisms; weaker (and requiring engagement with the ITPGRFA framework) for cultivated crop genetic resources. Informants consulted at SBSTTA-28/SBI-7 reinforced this nuance: ‘centre of origin/diversity’ and ‘documented use of that country’s material’ were ranked above raw biodiversity share as preferred disbursement factors (see Section 4.6), suggesting that Africa’s negotiating position should advocate for a multi-criteria disbursement formula rather than relying solely on biodiversity percentage.

### 2.5 The Provider-User Asymmetry

The DSI governance literature characterises the core inequity as a provider-user asymmetry: biodiversity-rich countries in the Global South provide the non-human genetic resources from which DSI is derived, while the commercial capacity to exploit that DSI is concentrated in high-income countries.[14,15] Scholz and colleagues (2022) documented that biodiverse nations, many of which are low- and middle-income countries, perceive their sovereign rights to have been undermined because potential monetary gains from the commercialisation of DSI are not shared back to them.[16]

Ljungqvist and colleagues (2025), in the most comprehensive cross-national ABS policy review, documented that as of May 2024, only 22 of 193 countries (11%) had any legislative reference to DSI, while 171 (89%) had no codified position. Critically, several of these 22 countries use terminology other than ‘DSI’, such as ‘genetic sequence data’ or ‘biological information’, indicating that ABS governance assessments restricted to the term ‘DSI’ will produce false negatives, as confirmed by the Namibia example raised at the side event (28 July 2026).[17]

## 3. Materials and Methods

### 3.1 Study Design

This study employs a structured evidence synthesis combining secondary data analysis with a structured expert elicitation. The secondary analysis follows PRISMA 2020 reporting elements where applicable; no formal risk-of-bias assessment was conducted, as this is not a clinical systematic review. The primary component is a structured key-informant elicitation of CBD delegates and experts at SBSTTA-28 and SBI-7 (Nairobi, July-August 2026). A PRISMA flow diagram is provided as supplementary Figure S1.

The search was restricted to January 2022 - April 2026 to focus on post-KMGBF and post-Decision 15/9 literature; earlier foundational legal instruments were included irrespective of date. A total of 412 records were identified before deduplication; 298 proceeded to title/abstract screening; 61 to full-text review; 34 peer-reviewed articles and 19 institutional documents met all inclusion criteria.

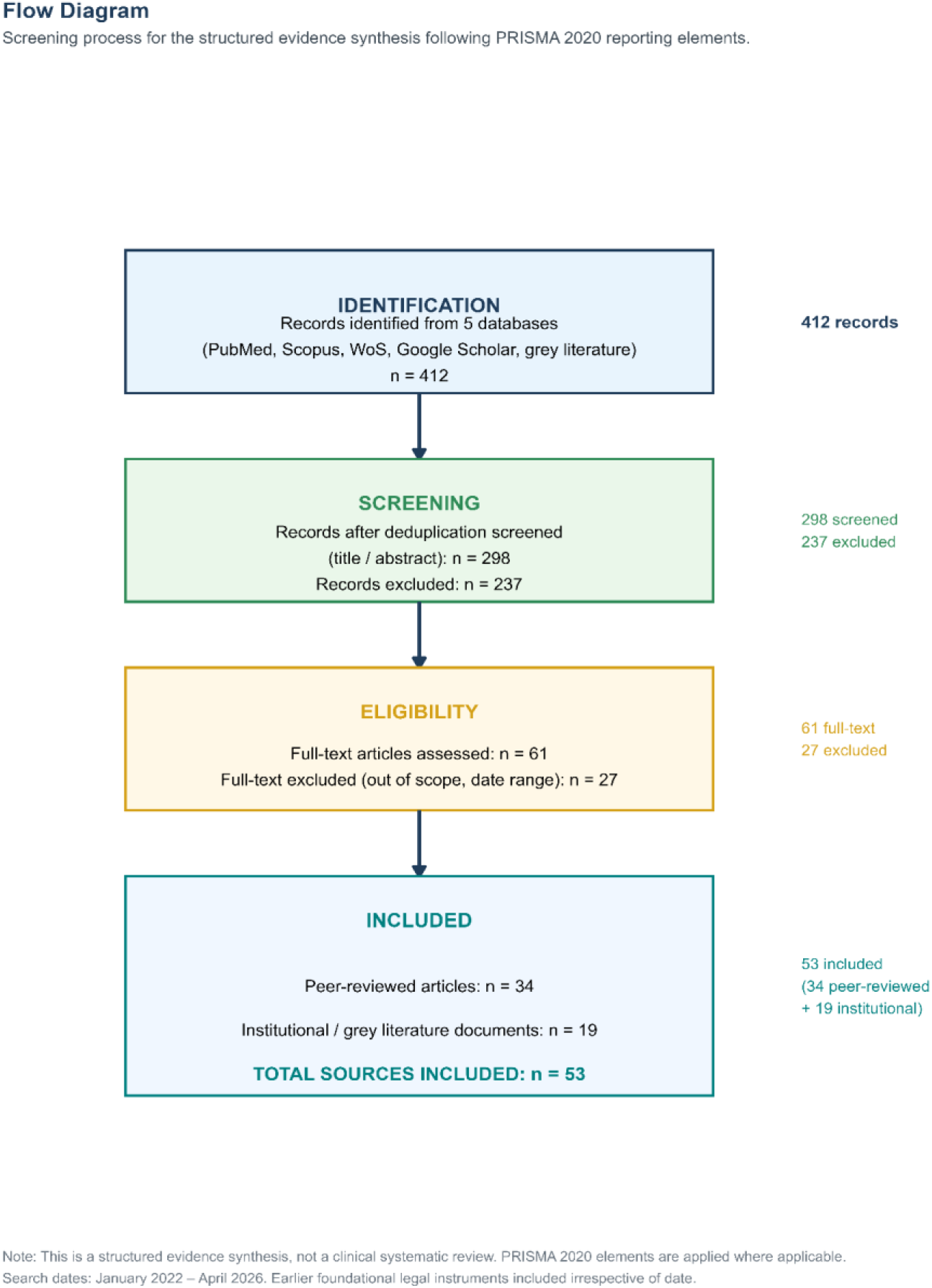

### 3.2 Literature Review

In response to methodological concerns raised at the DSI side event (28 July 2026), the search was expanded beyond the term “DSI” to include equivalent terminology used in national legislation and older literature. Search strings included: (1) (“digital sequence information” OR “DSI” OR “genetic sequence data” OR “biological sequence information” OR “nucleotide data”) AND (“Africa” OR “Sub-Saharan Africa”) AND (“benefit sharing” OR “access and benefit” OR “genetic resources”); (2) (“Nagoya Protocol” OR “ABS legislation” OR “access legislation” OR “bioprospecting law”) AND (“Africa” OR “developing countries”); (3) (“Cali Fund” OR “COP16 DSI” OR “multilateral mechanism DSI” OR “DSI fund”); (4) (“BBNJ” OR “WIPO genetic resources” OR “FAO ITPGRFA DSI” OR “WHO pathogen sharing”); (5) (“biodiversity hotspots Africa” OR “species richness Africa” OR “African endemic species”); (6) (“GWAS diversity” OR “genomic representation Africa”), as a secondary structural exclusion indicator only, not as a Cali Fund allocation basis. Inclusion criteria: peer-reviewed or authoritative institutional publication between January 2022 and April 2026; relevance to at least one study objective; availability in English or French.

### 3.3 ABS Governance Assessment with Expanded Search Criteria

The ABS governance readiness of each of the 54 AU member states was assessed using the CBD ABS Clearing-House (absch.cbd.int, accessed April 2026), the Nagoya Protocol ratification tracker (cbd.int/abs), and the policy database of Ljungqvist and colleagues (2025). Critically, assessment was not restricted to legislation explicitly using the term “DSI”. Country legislation using terms such as “genetic data,” “biological sequence information,” “biotechnology resources,” or “intangible components of genetic resources” was identified and coded separately as “pre-DSI equivalent coverage”, a new sub-category reflecting the Namibia-type examples raised at the side event. Countries were classified into five categories: (I) comprehensive ABS law with explicit non-human DSI coverage; (II) ABS law with equivalent pre-DSI terminology that may cover non-human DSI; (III) ABS law with ambiguous or partial coverage; (IV) Nagoya Protocol ratified but no domestic ABS implementation law; and (V) Nagoya Protocol not yet ratified. Classifications were cross-validated against the GIZ ABS Capacity Development Initiative database.

### 3.4 Quantitative Modelling

Revenue projections used the UNEP-WCMC/KPMG (2024) commissioned study (CBD/WGDSI/2/2/Add.2) as the primary baseline for non-human DSI-dependent sectors. These figures represent total sector revenues, not revenues attributable solely to DSI-enabled products; the UNEP-WCMC/KPMG methodology’s DSI attribution itself represents a source of uncertainty. No human genomics revenues are included. Africa’s modelled allocation share (20-25%) is the authors’ analytical construct based on Africa’s share of global non-human species richness and biodiversity hotspots; it is not contained in any agreed Cali Fund disbursement formula. The domesticated species qualification (Section 2.4) means this figure most accurately reflects Africa’s wild biodiversity allocation claim; the claim for domesticated crop DSI must be pursued separately through the ITPGRFA framework.

### 3.5 Structured Expert Elicitation: Key-Informant Design (SBSTTA-28/SBI-7, Nairobi, July - August 2026)

The primary component of this study is a structured expert elicitation, not a population survey. Expert elicitation is an established method for informing environmental and public-policy decisions where the relevant knowledge is concentrated in a small, identifiable community of specialists and where representative population data are not yet available.[45] Its sampling logic is purposive (key-informant) rather than probabilistic: informants are chosen deliberately because they hold the positional knowledge relevant to the question, and the criterion of adequacy is the quality and relevance of the informants’ expertise rather than the size of the sample.[46,47] Accordingly, the aim here is to map and triangulate the preferences of the actors who will shape the COP17 negotiation, not to estimate the distribution of opinion in any wider population.

The target population for this elicitation is intentionally narrow and bounded: accredited CBD Party delegates, IPLC representatives, scientific experts, and national ABS focal points actively engaged in the DSI agenda at the SBSTTA-28 and SBI-7 sessions. This is a small and hard-to-access expert community; the delegates negotiating DSI at these specific sessions number in the low hundreds, and the 25 informants consulted are drawn from precisely this group. Elicitation was administered on the margins of SBSTTA-28 (29-31 July 2026) and SBI-7 (1-4 August 2026). Participation was voluntary and anonymous, and responses are reported only in aggregate.

The instrument covered: (i) preferred Cali Fund contribution models; (ii) preferred disbursement factors; (iii) benefit-sharing modalities most valuable for Africa; (iv) relevance of other international processes; (v) ABS legislation status; (vi) governance-model preferences; and (vii) what Africa most needs to benefit fairly from DSI. Twenty-five complete responses were received from representatives of 14 countries: Brazil, Cameroon, Grenada, Kenya (multiple), Malawi, Malaysia, Mauritania, Nigeria, South Sudan, Sudan, United Kingdom, Zambia, Zimbabwe, and Antigua and Barbuda. Informant groups included Party delegates, IPLC representatives, NGO observers, scientists, and ABS focal points.

#### 3.5.1 Analytical treatment and rationale for adequacy

Results are analysed and reported as qualitative, count-based key-informant evidence. Findings are expressed as raw counts of informants (e.g., ‘20 of 25 informants’) rather than as inferential percentages, and no significance tests, confidence intervals, or population estimates are computed, because the purposive design does not support population inference.[46] This reporting choice is deliberate: presenting the data as expert testimony, rather than as rates generalised to a population, is both the honest and the methodologically correct treatment of a purposive sample.

The adequacy of 25 informants is judged against the standards of expert elicitation, not of probability sampling. In elicitation and key-informant designs, a smaller panel of well-placed experts is preferred over a larger, less specialised sample, and even modest panels drawn from the relevant expert community are considered informative when the informants collectively hold the knowledge in question.[45,46,47] Three features support adequacy here: (i) the informants occupy the exact decision-making roles the study concerns (Party delegates, ABS focal points, IPLC representatives); (ii) they span 14 countries and multiple stakeholder categories, giving the panel breadth across perspectives; and (iii) the purpose is hypothesis-generating and position-mapping ahead of COP17, not measurement of a population parameter. The elicitation is therefore best read as a structured scoping of expert opinion that *motivates and frames* the representative, field-based primary-data agenda set out in Section 7, and as one convergent strand of evidence alongside the literature synthesis rather than a stand-alone basis for the study’s conclusions.

Limitations of this design are stated plainly and revisited in Section 8. As a purposive, non-probability panel, the elicitation is subject to selection and self-selection effects and cannot be generalised to all African delegations, to IPLCs, or to global CBD Parties; its findings are indicative of expert and negotiator perspectives only. These limitations are precisely what the primary-research agenda in Section 7 (Gap 2) is designed to address.

## 4. Results

### 4.1 Africa’s Non-Human Biodiversity Endowment

#### 4.1.1 Terrestrial species richness and endemism

Africa, comprising 20% of the Earth’s land surface, harbours extraordinary non-human biological diversity. Bezeng and colleagues (2025) confirm that the continent hosts approximately one-quarter of the world’s mammal and bird species, with large fractions of global plant richness endemic to the continent.[18] A preprint systematic assessment (Sinsin et al., 2025, not peer-reviewed) documents approximately one-sixth of global plant richness, 17% of global mammal richness, 2,500 bird species, 950 amphibian species, and 5,000 freshwater fish species.[19] Only 19% of Africa’s landscape and 17% of its seascape are formally protected, well below the KMGBF 30×30 target.[18]

Nine of the world’s 36 formally recognised biodiversity hotspots fall wholly or substantially within African borders: the Cape Floristic Region, Succulent Karoo, Maputaland-Pondoland-Albany, Coastal Forests of Eastern Africa, Eastern Afromontane, Guinean Forests of West Africa, Horn of Africa, Madagascar and the Indian Ocean Islands, and the Mediterranean Basin (partially).[3,20] Africa’s 25% share of global biodiversity hotspots (9/36) is the primary quantitative basis for the upper bound of its modelled Cali Fund allocation share, with the domesticated species qualification noted in Section 2.4.

The Congo Basin warrants specific attention as a non-human DSI resource. Covering approximately 240 million hectares across eight African countries, it is among the least genomically characterised ecosystems on Earth and harbours tens of thousands of plant and animal species whose sequences have not been deposited in any public database. Its extensive unsequenced diversity represents both a governance gap and a non-monetary benefit-sharing opportunity: prioritising African bioinformatics infrastructure would enable sequencing of this irreplaceable resource.

**Table 2.** Africa’s Share of Global Non-Human Biodiversity.

| Non-Human Biodiversity Metric | Africa (estimate) | Global Total | Africa Share |
| --- | --- | --- | --- |
| Land area (km<sup>2</sup>) | 30.4 million | 149 million | 20% |
| Terrestrial species richness | ≈25% of global species | All species | ~25% |
| Biodiversity hotspots | 9 hotspots | 36 globally | 25% |
| Mammal species | ≈1,100 | ≈5,500 | ~20% |
| Bird species | ≈2,500 | ≈10,000 | ~25% |
| Amphibian species | ≈950 | ≈8,000 | ~12% |
| Plant species | ≈50,000 | ≈390,000 | ~13% |
| Protected terrestrial area | 19% | 30% (KMGBF target) | Below target |

*Note.* KMGBF = Kunming-Montreal Global Biodiversity Framework; ≈ = approximately.
*Sources:*
[3,18,19,20]. Sinsin et al. (2025) [19] is a preprint, not peer reviewed. Africa's wild species richness and hotspot share (25%) is the primary basis for the upper bound of the modelled Cali Fund allocation share for wild biodiversity. Domesticated crop DSI requires separate analysis through the FAO ITPGRFA framework.

#### 4.1.2 Human genomic diversity: contextual indicator only, outside CBD/Cali Fund scope

Africa’s status as the evolutionary cradle of anatomically modern Homo sapiens, as dated by the Jebel Irhoud fossil record (Hublin et al., 2017), confers genetic diversity that is scientifically without parallel.[21] The African Genome Variation Project (Gurdasani et al., 2015) sequenced 320 individuals from 18 African populations and identified over three million novel variants not previously described.[22] The AGenDA project (Ramsay et al., 2026) generated whole-genome sequences from over 1,000 individuals across nine countries (Angola, Democratic Republic of Congo, Kenya, Libya, Mauritius, Rwanda, South Africa, Tunisia, Zimbabwe).[23]

Corpas and colleagues (2025) documented that 94.48% of samples in the GWAS Diversity Monitor were of European ancestry as of September 2024, while Sub-Saharan African ancestry samples constituted approximately 0.47% of the GWAS Catalog as of 2023, having declined from approximately 1.1% in 2019.[24] This underrepresentation is presented here as evidence of the same structural exclusion, infrastructure deficits and investment asymmetries, that simultaneously marginalises Africa in non-human DSI databases. It is not part of the Cali Fund allocation argument.

### 4.2 Africa’s Underrepresentation in Non-Human DSI Databases: Collection, Attribution, and Governance Gaps

A key question raised at the side event (28 July 2026) is whether Africa’s absence from global DSI databases reflects a collection problem (African biodiversity is not being sequenced), a metadata attribution problem (African organisms are being sequenced but not attributed to their country of origin), or both. Evidence suggests both mechanisms operate simultaneously.

On the collection side: the African BioGenome Project (AfricaBP), formally launched in 2021 with the objective of sequencing 105,000 African species, has documented that the vast majority of Africa’s known biological diversity remains unsequenced.[25,26] The Genomic Reference Resource for African Cattle (GRRFAC) illustrates the concrete scale: despite Africa hosting over 150 indigenous cattle breeds adapted over millennia, fewer than 300 individuals had reference-quality genome sequences prior to the GRRFAC initiative.[27]

On the metadata attribution side: billions of legacy records in GenBank and the INSDC carry no country-of-origin metadata. The 2023 INSDC spatiotemporal annotation update addresses this for future submissions but not for the existing archive.[2] The side event participants recommended that the research team engage directly with GenBank, ENA, and DDBJ database curators to understand what provenance metadata exists in internal records but is not publicly displayed, the data ‘sits somewhere.’

### 4.3 Commercial Non-Human DSI Utilisation: Sector Revenue Analysis

The UNEP-WCMC/KPMG (2024) commissioned study provides the most authoritative published estimate of revenues generated by non-human DSI-dependent sectors. These figures represent total sector revenues, not revenues attributable solely to DSI-enabled products.[1]

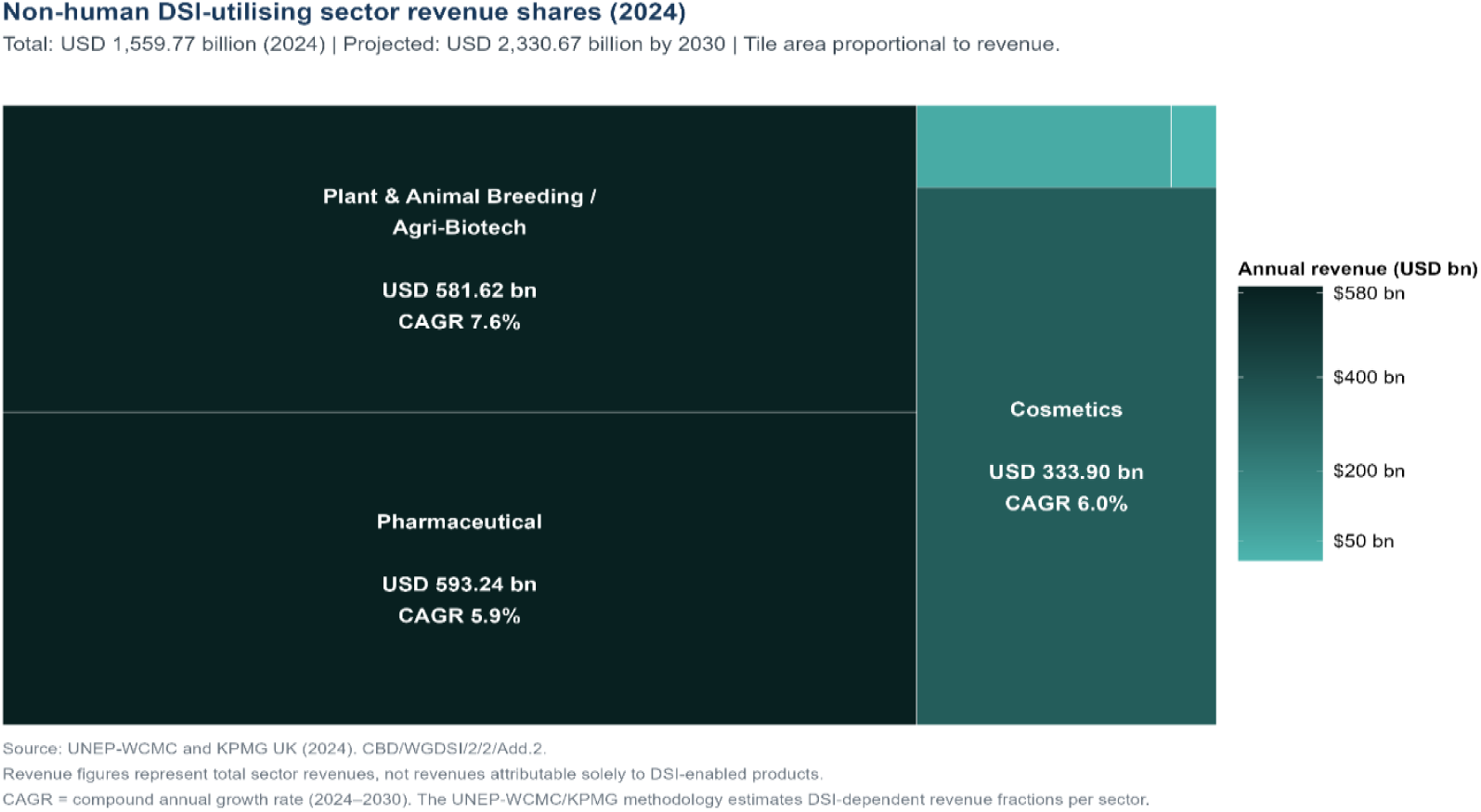

### 4.4 ABS Governance Landscape: Systematic Assessment with Expanded Classification

#### 4.4.1 Continental overview, addressing Africa’s national heterogeneity

Systematic assessment of the ABS legislative landscape across all 54 AU member states, using an expanded classification that includes pre-DSI equivalent genetic resources legislation, reveals significant national heterogeneity within the broadly fragmented continental picture.[17,28]

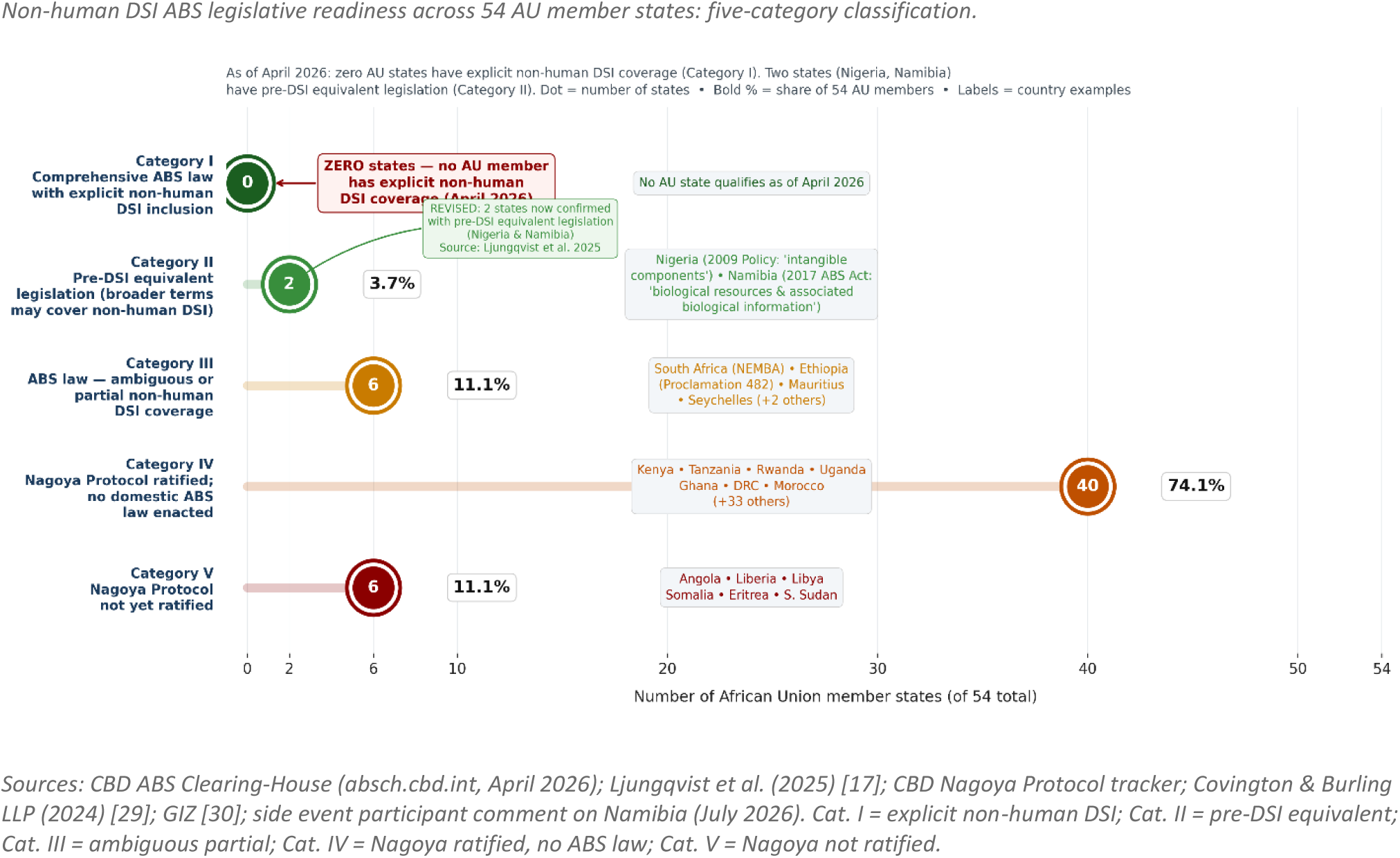

#### 4.4.2 The three most critical structural barriers to non-human DSI governance

Rather than enumerating barriers at equal weight, three structurally critical impediments are identified.

The most fundamental barrier is database traceability and the collection gap. Africa’s absence from INSDC databases reflects both a collection gap (biodiversity not being sequenced at scale) and a metadata attribution problem (sequences deposited without country-of-origin metadata). These are legally distinct problems: the collection gap requires investment in African sequencing infrastructure; the attribution gap requires engagement with database curators and the INSDC’s 2023 spatiotemporal annotation policy.[2]

The second barrier is the ‘public domain’ legal narrative. Many courts and legal commentators treat unmodified DSI as unpatentable public domain information, with IP rights arising only from downstream ‘innovation.’ This narrative is a legal distortion: it ignores that African environments, shaped over centuries by indigenous selection, management, and modification (such as African dark earths and centuries of traditional crop breeding), have created the genomic substrates that bioprospectors seek. The WIPO GR Treaty’s patent disclosure requirement creates a new tool to challenge this narrative by requiring applicants to reveal the source of genetic resources in patents.[10]

The third barrier is voluntary mechanism design and enforcement absence. Decision 16/2 establishes voluntary, indicative contribution rates. In the expert elicitation (Section 4.6), 20 of the 25 informants favoured mandatory mechanisms. The IFPMA’s public resistance after COP16 illustrates the risk of companies opting out.[31] Africa’s negotiating position at COP17 should prioritise transforming indicative rates into mandatory obligations with a compliance mechanism, as modelled by the WHO PIP Framework’s benefit-sharing structure for influenza viruses.[11]

### 4.5 Cali Fund: Revenue Modelling and Africa’s Modelled Allocation Share

Using the UNEP-WCMC/KPMG (2024) non-human DSI sector baseline and applying Decision 16/2’s indicative rates, Table 5 presents projected annual Cali Fund revenues under five scenarios. Africa’s modelled allocation share (20-25%) is the authors’ analytical construct based on wild biodiversity metrics; it is not an agreed formula. Both weights are shown for 2024 and 2030 to make this explicit.[1]

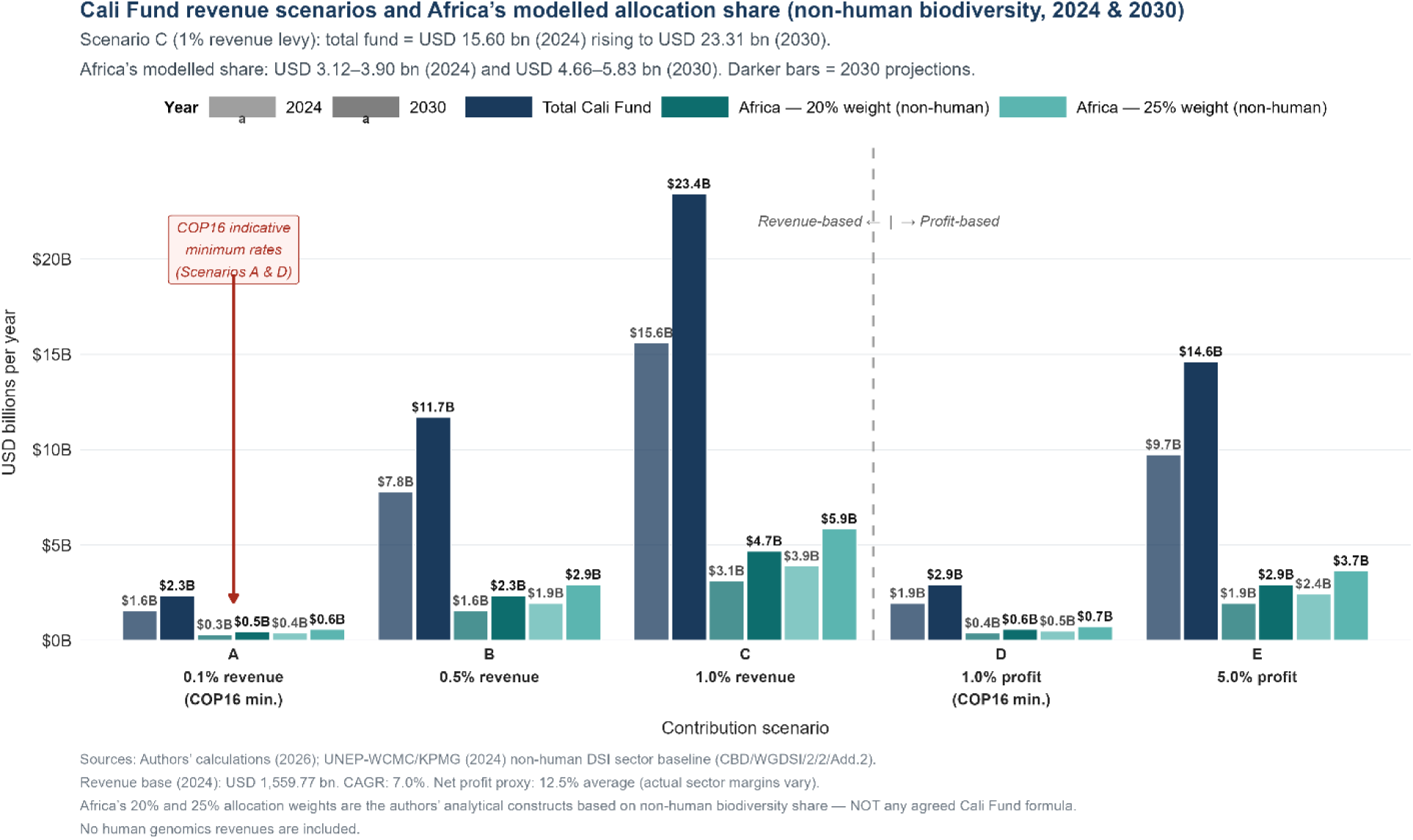

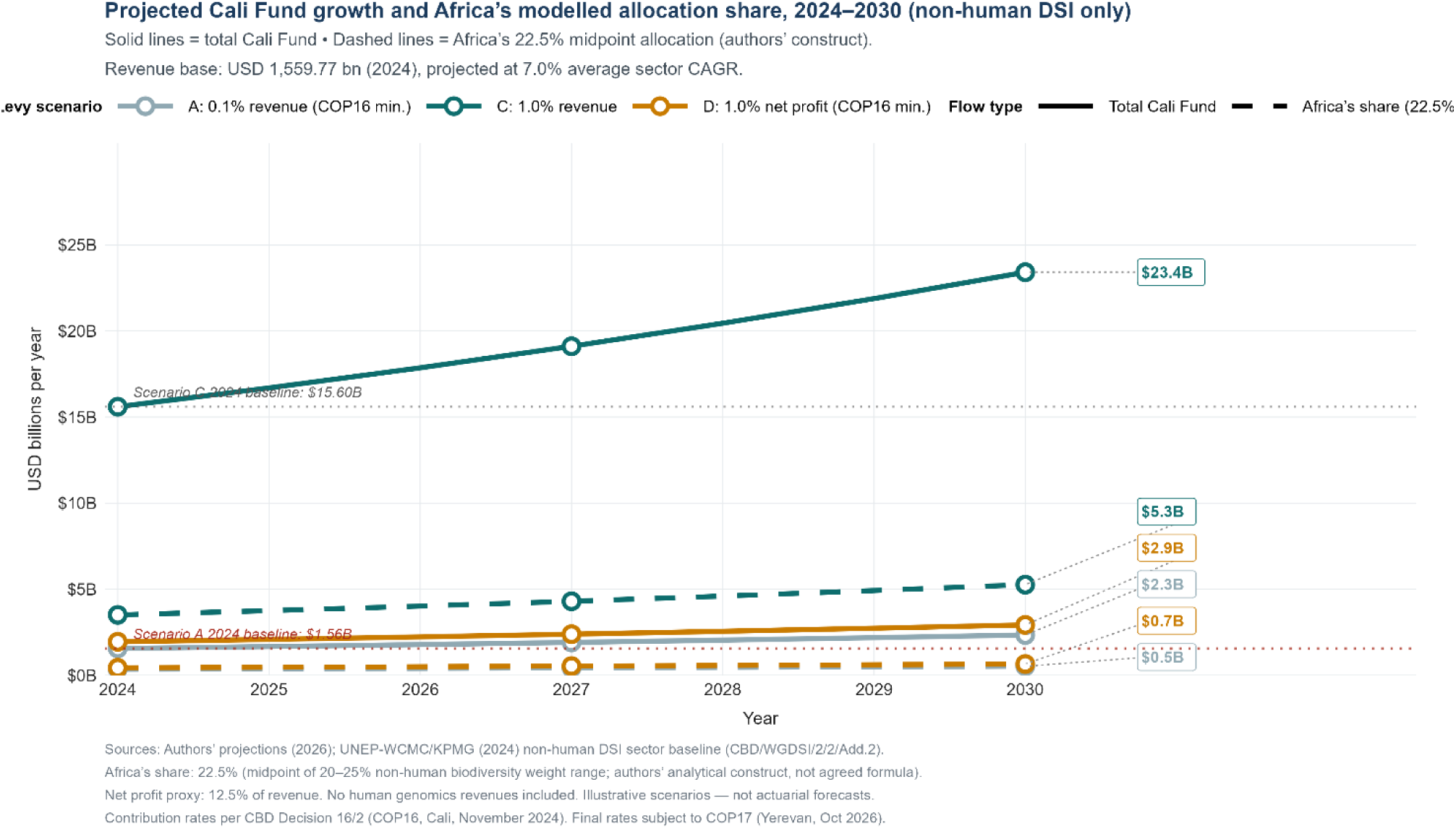

### 4.6 Expert Elicitation Results: Key-Informant Preferences at SBSTTA-28/SBI-7

This section presents the results of the structured expert elicitation of 25 CBD delegates and experts at SBSTTA-28 and SBI-7 (Nairobi, July-August 2026). Findings are reported as counts of informants and are *indicative of expert and negotiator perspectives; they are not statistically representative of global or African positions*, and they function throughout as corroboration for the literature synthesis rather than as independent proof (see Section 3.5).

**Table 3.** Key Findings from the Expert Elicitation and Their Implications for the Analysis.

| Elicitation Item | Key Finding (n=25 informants) | Implication for the Analysis |
| --- | --- | --- |
| Preferred Cali Fund contribution model | Mandatory (all or threshold): 20 of 25.<br>Voluntary: 4 of 25. | Corroborates the analysis's critique of the voluntary model. Africa's case for mandatory mechanisms at COP17 is reinforced by negotiator sentiment. |
| Most important disbursement factors | Centre of origin/diversity and documented use of a country's material ranked highest; raw biodiversity share ranked third. | Qualifies the 20–25% biodiversity-share model as a sole basis; supports a multi-criteria formula. |
| Domesticated species benefit-sharing (to whom?) | Informants split: multilateral pool; centre of origin; current host country. | Corroborates the reviewer's challenge. No consensus; Africa must engage FAO ITPGRFA. |
| Most valuable benefit-sharing forms | Capacity building and technology transfer ranked alongside or above monetary payments in 18 of 25 responses. | Non-monetary benefits should receive equal prominence in policy recommendations. |
| Other international processes (multi-select) | FAO ITPGRFA: 22 of 25; WHO/PIP: 20 of 25; BBNJ: 19 of 25; WIPO: 18 of 25. Only 2 of 25 said CBD sufficient alone. | Corroborates the multi-forum expansion of the analysis's scope. |
| ABS legislation covering non-human DSI | Explicit DSI: 1; general/partial: 8; none: 6; don't know: 6; in development: 3. | Consistent with the zero-explicit-coverage finding, and surfaces the pre-DSI equivalent category (8 with partial/general coverage). |
| Preferred governance model | Hybrid (CBD + other forums): 10 of 25; multilateral (CBD only): 8 of 25; regional African: 5 of 25. | Hybrid and multilateral preferred; consistent with multi-forum framing. |
| What Africa most needs for DSI | Infrastructure and metadata standards; funding; capacity building; legal support. | Policy recommendations should lead with non-monetary, infrastructure-first priorities. |

Elicitation conducted on the margins of SBSTTA-28 (29–31 July 2026) and SBI-7 (1–4 August 2026), Nairobi, Kenya. Purposive key-informant panel; not statistically representative. Informants from: Brazil, Cameroon, Grenada, Kenya (multiple), Malawi, Malaysia, Mauritania, Nigeria, South Sudan, Sudan, United Kingdom, Zambia, Zimbabwe, and Antigua and Barbuda. Informant groups: Party delegates, IPLC representatives, NGO observers, scientific experts, ABS focal points.

## 5. Discussion

### 5.1 The Multi-Forum DSI Governance Architecture and Africa’s Strategic Response

The most structurally significant finding of this study, established through literature analysis and corroborated by the expert elicitation, is that DSI governance is not a single-forum matter. Among the 25 informants, 22 identified FAO ITPGRFA as a relevant parallel process; 20 identified the WHO framework; 19 identified BBNJ; and 18 identified WIPO. Only 2 informants considered the CBD Cali Fund mechanism sufficient on its own. While these counts describe the panel rather than a wider population, they align closely with the documentary evidence on the distribution of DSI governance across forums.

This has direct implications for Africa’s COP17 strategy. A continental DSI governance approach restricted to CBD negotiations will miss the majority of commercially valuable DSI. Africa’s agricultural ministries must simultaneously engage ITPGRFA reform for crop DSI; health ministries must engage the WHO Pandemic Instrument negotiations for pathogen sequences; and trade ministries must engage WIPO implementation for patent-linked disclosure. The Agenda 2063 framework, specifically its aspiration for a prosperous Africa based on inclusive growth and sustainable development, provides the continental political mandate for this multi-forum engagement.

The BBNJ Agreement’s provenance-tracking model is particularly relevant for African terrestrial DSI governance: attaching unique identifiers to biological samples at collection would enable sequence records to be traced back to their country of origin even after database deposition. The side event (28 July 2026) recommended evaluating whether this identifier model, already negotiated for high-seas marine samples, can be adapted as a voluntary or mandatory standard for terrestrial African biodiversity programmes.[9]

### 5.2 Rethinking the Allocation Model: Beyond Biodiversity Share

The observation that 25% of global biodiversity does not automatically translate to 25% of Cali Fund disbursements is well-founded, and it was reinforced by the elicitation. The 20 - 25% biodiversity-share model in this analysis is most defensible for wild, endemic African species and ecosystems. For domesticated crops and livestock, whose genetic centres of origin often lie in Mesoamerica, the Fertile Crescent, or South Asia, Africa’s claim requires engagement with the FAO ITPGRFA multilateral system, not the CBD Cali Fund.

Informants consistently ranked ‘centre of origin/diversity’ and ‘documented use of that country’s material’ above raw biodiversity share as preferred disbursement factors. Africa’s negotiating position should therefore advocate for a multi-criteria disbursement formula that combines: (i) biodiversity hotspot density and wild species richness; (ii) documented country-of-origin metadata in INSDC databases; (iii) demonstrated conservation stewardship; and (iv) development status as a corrective factor.

### 5.3 Challenging the Voluntary Model: Evidence for Mandatory Mechanisms

The current Cali Fund model relies on voluntary, indicative contribution rates. The elicitation, in which 20 of 25 informants favoured mandatory mechanisms, offers direct qualitative evidence that the voluntary model has weak legitimacy among the very negotiators and experts who will shape COP17. This expert signal converges with documentary evidence, including the IFPMA’s public resistance after COP16, which confirms the practical risk of opt-out. Because the elicitation is purposive, this convergence, rather than the counts alone, carries the argument.

The WHO PIP Framework for pandemic influenza viruses offers an instructive model: it creates binding benefit-sharing agreements as a condition of access to the WHO’s global influenza surveillance network, generating both monetary contributions (to the PIP fund) and non-monetary benefits (technology transfer for vaccine manufacturing in developing countries). A similar structure, contributions as a condition of access to INSDC databases, could be explored for the Cali Fund’s long-term architecture.

### 5.4 Community Agency, IPLC Rights, and Data Sovereignty

A recurring theme across all three comment sources is that the IPLC dimension of DSI governance must go beyond positioning communities as recipients of Cali Fund disbursements. Indigenous peoples and local communities have their own biotechnologists, traditional breeders, and knowledge systems. Any benefit-sharing framework must recognise this existing scientific capacity and build governance mechanisms that give IPLCs decision-making power, not just access to project funding.

AUDA-NEPAD’s question, ‘Is Africa going to have sovereignty over the data?’, points to a dimension that the current Cali Fund architecture does not adequately address. Monetary benefit-sharing without data sovereignty may generate revenue flows while perpetuating a governance structure in which the sequencing infrastructure, database hosting, and data governance authority remain outside Africa. A more complete DSI governance framework for Africa must address: which institutions host African sequence data; who controls access; and how African researchers and communities are empowered as data producers rather than just data sources.

### 5.5 What Africa Most Needs: Infrastructure Before Revenue

Informants’ top priorities, infrastructure, metadata standards, capacity building, and legal support, align with the side event’s emphasis on non-monetary benefits. Even if the Cali Fund generates significant revenue flows, Africa cannot claim its share without: institutional infrastructure to access disbursements; bioinformatics capacity to generate and deposit sequences; metadata systems to document country-of-origin; and legal capacity to litigate against biopiracy. These infrastructure investments are prerequisites, not alternatives, to monetary benefit-sharing.

The analysis’s policy implications must therefore lead with non-monetary, infrastructure-first priorities: African community biobanks, national sequencing facilities, bioinformatics training programmes, and metadata standards adoption. Alignment with Agenda 2063’s aspirations for prosperity, institutional capacity, and science-driven development, and with the Malabo and Kampala Declarations on agricultural and biodiversity governance, provides the continental political mandate.

## 6. Policy Implications for African Institutions

Incorporating the evidence from this study and the perspectives elicited from 25 key informants at SBSTTA-28/SBI-7, eight priority action areas are identified for African institutions before COP17.

### 1. Strengthen ABS legislation with explicit non-human DSI coverage and pre-DSI equivalent recognition

No AU state has enacted legislation explicitly and comprehensively covering non-human DSI. Priority actions: (a) update existing ABS laws to include explicit non-human DSI definitions; (b) review existing genetic resources legislation for pre-DSI equivalent coverage (the Namibia model); (c) adopt harmonised non-human DSI provisions at REC level; and (d) formally incorporate the ‘old DSI vs new DSI’ distinction into national legislative frameworks.

### 2. Engage all relevant international forums, not just the CBD

Agriculture ministries must engage FAO ITPGRFA negotiations for crop and livestock DSI; health ministries must engage the WHO Pandemic Instrument for pathogen sequences; trade ministries must implement the WIPO GR Treaty patent disclosure requirement domestically. A national inter-ministerial coordination mechanism for DSI governance is urgently needed in each AU state.

### 3. Advocate for mandatory contribution mechanisms at COP17

The voluntary model is insufficient. Africa’s negotiating block should propose: mandatory minimum contributions as a condition of INSDC database access; a compliance verification mechanism; and penalties or access restrictions for persistent non-compliance. The WHO PIP Framework provides a negotiated precedent.

### 4. Invest in provenance infrastructure and database metadata standards

Africa must actively engage INSDC database curators (GenBank, ENA, DDBJ) to understand what provenance data exists in internal records. National sequencing programmes should adopt standardised collection-origin metadata as a prerequisite for deposition. The BBNJ sample identifier model should be evaluated for adoption in terrestrial biodiversity programmes.

### 5. Ensure IPLC agency, not just access

Decision 16/2’s 50% IPLC allocation is an opportunity and a risk. Without domestically enforceable community benefit-sharing protocols, funds may be absorbed at government level. African governments must: develop enforceable community benefit-sharing frameworks; establish direct IPLC access pathways to Cali Fund disbursements; and recognise existing IPLC scientific capacity in governance structures.

### 6. Define and demand meaningful non-monetary benefits

Generic capacity building commitments are insufficient. Africa must demand: technology transfer agreements for bioinformatics infrastructure; joint ownership of sequence data generated from African biodiversity; co-authorship requirements in benefit-sharing agreements; community-controlled biobanks; and inclusion of African institutions in product development processes that use African DSI.

### 7. Align with Agenda 2063 and continental political declarations

This research must be explicitly framed within the seven aspirations of Agenda 2063 (an integrated, prosperous, and peaceful Africa driven by its own citizens), the Malabo Declaration on agricultural growth and food security, and the Kampala Declaration on biodiversity and African heritage. DSI governance is not a standalone biodiversity issue; it is a dimension of Africa’s sovereign right to benefit from its own natural endowments.

### 8. Establish ethical and data sovereignty safeguards for research

Any primary data collection on African genetic resources or traditional knowledge must include robust ethical safeguards: community consent protocols; data sovereignty agreements specifying that African data is hosted on African-controlled infrastructure; transparent beneficiary mapping showing how research benefits flow to communities; and protections against research output becoming a bioprospecting tool. These safeguards must be in place before field data collection begins.

### 7. Research Gap and Future Directions

This study is a structured evidence synthesis whose primary component is a purposive expert elicitation. Its findings are robust within the limits of secondary data and key-informant evidence, but five critical knowledge gaps constrain the conclusions that can be drawn and point directly to the representative primary-research agenda needed. Gap 2 in particular is the direct counterpart to the elicitation’s design: it specifies the representative, field-based data collection that would move beyond the indicative, purposive panel reported here.

#### Gap 1: Systematic country-of-origin metadata extraction from INSDC databases

No published study provides a systematic, quantitative account of how many sequences in GenBank, ENA, or DDBJ originate from African countries, by taxonomic group and sub-national geography. The conclusion that Africa is severely underrepresented is well-supported by proxy indicators (AfricaBP data, GRRFAC, INSDC own disclosures) but cannot be precisely quantified without a systematic metadata extraction study. Future researchers should: (i) use the INSDC API to extract all records with collection location fields; (ii) map the proportion of sequences traceable to African countries by country, taxonomic group, and decade; (iii) distinguish records with missing metadata from those with confirmed non-African origin; and (iv) engage GenBank, ENA, and DDBJ curators directly about what provenance data exists in internal records not publicly displayed.

#### Gap 2: Representative, field-based primary data from IPLCs and delegations

The expert elicitation reported here is a purposive key-informant exercise and, by design, cannot be generalised to all African delegations, to IPLCs, or to the wider body of CBD Parties. This study therefore has no representative primary data from indigenous peoples and local communities in Africa about their awareness of the Cali Fund, their expectations of benefit-sharing, or their existing governance mechanisms for genetic resources. Future research should convert the indicative findings of this elicitation into a representative evidence base through: (i) a probability-based or stratified survey of accredited CBD delegations across AU member states with a pre-registered sampling frame and reported response rates; and (ii) semi-structured interviews and participatory assessments in at least 8-10 African countries, with communities representing diverse ecological zones, language families, and governance traditions. Particular attention should be given to communities whose territories overlap with biodiversity hotspots. The elicitation instrument and item structure used here can serve as the tested basis for such an expanded, representative study.

#### Gap 3: Corporate DSI supply chain tracing

The UNEP-WCMC/KPMG revenue figures represent total sector revenues, not revenues demonstrably attributable to African DSI. A supply chain analysis tracing a selection of pharmaceutical, cosmetic, and agricultural patents to their genetic resource origins, using the WIPO GR Treaty’s new patent disclosure database when it becomes operational, would provide the first empirical quantification of Africa’s contribution to specific commercial products. This would transform the allocation model from a proxy estimate to an evidence-based claim.

#### Gap 4: Comparative legal analysis of pre-DSI national frameworks

This study has identified the category of ‘pre-DSI equivalent legislation’ (exemplified by Namibia’s 2017 ABS Act) but has not systematically mapped which AU states have such frameworks or assessed their legal effectiveness in extending to non-human DSI. A systematic comparative legal analysis across 54 AU states, using search terms beyond ‘DSI’ and consulting in-country legal experts, would substantially refine the governance readiness assessment and identify immediate legislative entry points that do not require new laws.

#### Gap 5: Multi-forum coordination and overlap mapping

This study has described the multi-forum DSI governance architecture at a conceptual level. A practical mapping of which specific categories of DSI relevant to African biodiversity fall under each instrument, and where the current instruments create governance gaps, overlaps, or contradictions, would provide the operational foundation for Africa’s multi-forum engagement strategy. This would require input from legal experts in CBD, ITPGRFA, WHO, BBNJ, and WIPO processes simultaneously.

Future research addressing these five gaps would transform this study from a governance analysis into a quantified evidence base for Africa’s future negotiating position. The combination of corporate supply chain tracing (Gap 3), representative community-level and delegation field data (Gap 2), and precise database extraction (Gap 1) would constitute the most comprehensive primary evidence base on African DSI governance yet assembled.

## 8. Conclusions

This study presents six evidence-supported conclusions.

First, Africa is among the most biologically rich continents on Earth for non-human life, with 9 of 36 global biodiversity hotspots and approximately 25% of terrestrial species richness. This wild biodiversity endowment is the primary scientific basis for Africa’s modelled Cali Fund allocation claim, with the important qualification that domesticated crop and livestock DSI requires separate engagement through the FAO ITPGRFA framework.

Second, DSI governance is a multi-forum matter. The CBD Cali Fund is the primary mechanism for wild biodiversity DSI, but FAO ITPGRFA, the WHO Pandemic Instrument, the BBNJ Agreement, and the WIPO GR Treaty each govern distinct DSI categories relevant to Africa. A CBD-only strategy is insufficient. The expert elicitation was consistent with this: only 2 of 25 informants considered the CBD mechanism sufficient on its own.

Third, Africa is profoundly underrepresented in non-human DSI databases through both a collection gap and a metadata attribution problem. Legacy data traceability, the ‘public domain’ legal narrative, and the voluntary mechanism design are the three most critical structural barriers. The WIPO GR Treaty’s patent disclosure requirement and the BBNJ provenance identifier model offer new tools for addressing these barriers.

Fourth, ABS governance readiness is fragmented and heterogeneous across 54 AU states. Zero states have legislation explicitly covering non-human DSI, but several (including Namibia) have pre-DSI equivalent frameworks that may provide partial coverage. The expanded five-category classification introduced in this study provides a more accurate governance baseline than previous binary assessments.

Fifth, the voluntary contribution model has weak legitimacy among the negotiators and experts consulted: 20 of 25 key informants favoured mandatory mechanisms, a signal that converges with the documentary evidence. Africa’s COP17 position should prioritise transforming indicative voluntary rates into mandatory obligations with a compliance mechanism, drawing on the WHO PIP Framework as a precedent.

Sixth, realising Africa’s modelled allocation share requires infrastructure investment before revenue capture, multi-forum governance engagement, mandatory contribution mechanisms, and robust IPLC agency frameworks. Because the primary evidence here is a purposive expert elicitation, the research gaps identified in Section 7, and Gap 2 in particular, define the representative primary-data agenda that will convert these indicative findings into a conclusive evidence base ahead of future CBD review processes.

## 9. Declarations

### Data availability

Elicitation data, datasets, and R scripts used for quantitative analysis are available from the corresponding author on reasonable request. All primary data sources are cited in the reference list.

### Author contributions (CRediT)

Yves Shema: Conceptualization, Methodology, Formal analysis, Software, Visualization, Writing; original draft. Suwilanji Sinyangwe: Conceptualization, Writing; original draft, Writing; review & editing, Project administration. Fredrick Ayodele: Conceptualization, Writing – original draft, Methodology, Validation.

## Acknowledgements

The authors thank the delegates and experts who participated in the DSI Governance and Benefit-Sharing in Sub-Saharan Africa side event (28 July 2026) and the SBSTTA-28/SBI-7 expert elicitation (Nairobi, July-August 2026) for their invaluable insights. Special thanks to Pierre du Plessis (SADC) and Mphatso Martha Kalemba (Malawi Ministry of Natural Resources and Climate Change for their substantive comments on the earlier version of this paper, and to the African Wildlife Foundation for financial and logistical support.

